# Raptor-informed feathered drone reveals tail-twist functions in avian turning manoeuvres

**DOI:** 10.1101/2023.12.29.573672

**Authors:** Hoang-Vu Phan, Dario Floreano

## Abstract

Banked turn is a common flight manoeuvre observed in birds and aircraft. To initiate the turn, whereas traditional aircraft rely on the wing ailerons, most birds use a variety of asymmetric wing morphing control techniques, validated in engineered replicas, to roll their bodies and thus redirect the lift vector to the direction of the turn. Nevertheless, when searching for prey, soaring raptors execute steady banked turns without exhibiting observable wing movements apart from tail twisting around the body axis. Despite the role as a vertical stabilizer in traditional aircraft, the reasons why birds twist the tails in banked turn are still not well understood. Here, we use an avian-inspired feathered drone to find that the tail located in proximal arrangement behind the wings enters wing-induced asymmetric flow region during twisting and generates asymmetric lift that results in both roll and yaw moments sufficient to coordinate banked turns. Moreover, twisting the tail induces a nose-up pitch moment that increases the angle of attack of the wings, thereby generating more lift that compensates for losses caused by the banking motion. Flight experiments confirm the effectiveness of tail twist to control not only steady low-speed banked turns but also high-speed sharp turns by means of coordinated tail twist and pitch with asymmetric wing shape morphing. These findings contribute to the understanding of avian flight behaviours that are difficult to study in controlled laboratory settings, and provide effective control strategies for agile drones with morphing aerial surfaces.

**One sentence summary:** Raptor-informed feathered drone reveals that twisting the tail located at the trailing edges of the wings generates aerodynamic control forces caused by wing-induced asymmetric flow to let birds execute both steady banked turns and high-speed sharp turns.

## Introduction

Soaring raptors including eagles, hawks, and falcons, are frequently observed twisting their tail around the body axis during banked turns^1,2^, which are common flight manoeuvres. In the entry phase of a banked turn, birds twist the tail in the opposite direction to the bank, and in the exit phase reverse the tail rotation^1,3^. It is well known that birds twist the tail during turns to compensate for adverse yaw generated by asymmetric drag on the wings^1,2,4–6^. However, the mechanism that initiates the roll moment to bank the bird’s body is still not well understood. Theoretical studies that model bird tails as delta wings predict that twisting the tail around the body axis at high angles of attack induces asymmetric leading-edge vortices due to sideslip and, consequently, generates both roll and yaw moments sufficient to initiate a banked turn^7–9^. However, these studies assume uniform flow and vortices that remain attached to the wing upper surface even at high angles of attack; furthermore, these models do not consider potential airflow disruptions over the tail generated by the proximity of the bird’s wings and body^9–11^. Wind tunnel experiments with simplified fixed-wing models and rotating flat-plate tails^6,12^ reported the generation of roll moment by tail twisting, but the magnitude was too small for effective roll control in birds^4,13^. Understanding the aerodynamic function of tail twist in avian turning manoeuvres thereby necessitates consideration of the full flight envelope, but obtaining such flight behavior with birds in controlled laboratory settings is nontrivial.

## Results

### Raptor-informed feathered drone

Here, we study the avian turning manoeuvres by means of a raptor-informed feathered drone with morphing wings and tail (Fig. 1, ‘Feathered wing and tail design’ in Methods, Extended Data Figs. 1-3, and Extended Data Table 1). The wing skeleton consists of three segmented linkages replicating the bird’s humerus, ulna and manus that respectively rotate around the shoulder, elbow, and wrist joints with dimensions and mass similar to those of raptor wings^14,15^ (Fig. 1c,d, and Extended Data Fig. 1). Birds can couple their elbow and wrist motion to change the shape and area of the wing^16,17^. We also coupled the motion of the shoulder, elbow and wrist joints by a single tendon-driven actuator (Extended Data Fig. 1). Bird wings and tail are fully covered by lightweight overlapping feathers^18^, which we approximated by means of folding foam structures (Fig. 1d, ‘Feathered wing and tail design’ in Methods, and Extended Data Figs. 1-3) that allow the drone to reduce wing length, wing surface and tail surface by approximately 60%, 53% and 68%, respectively, when fully folded (Fig. 1b).

**Fig. 1.**
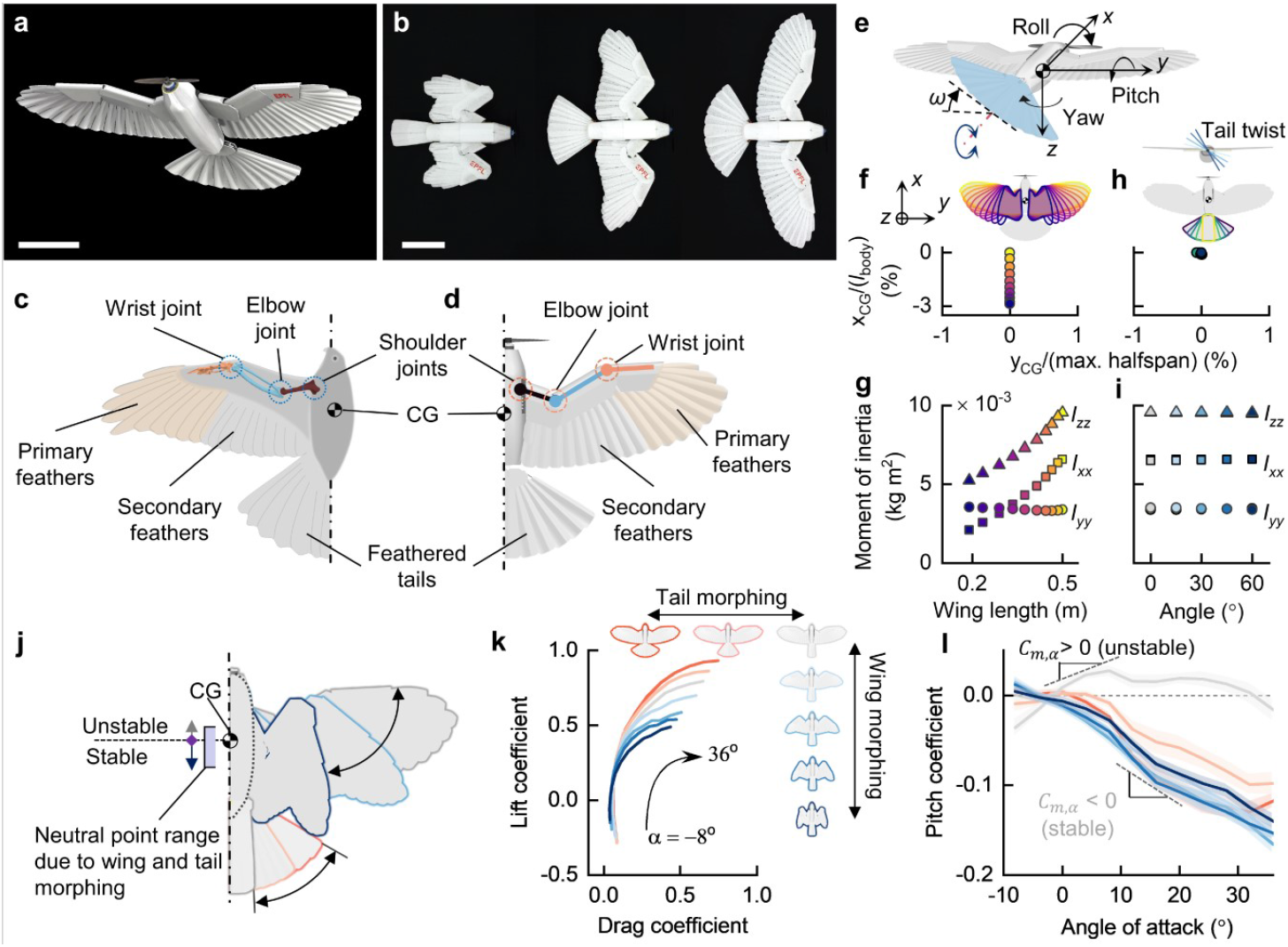
Bio-informed design of feathered morphing drone. **a**, The feathered drone with avian-inspired morphing wing and tail. **b**, Shape morphing allows the drone to change the spans and surface areas of the wings and tail. Scale bar: 20 cm. **c**,**d**, Coupled motions of bird wing skeletons around elbow and wrist joints (**c**) inspires design of the robotic wing skeleton (**d**). **e**, Bird-inspired rudderless tail capable of twisting around the body axis. **f**,**g**, Wing shape morphing minimally alters location of the CG (**f**) but strongly affects roll and yaw moments of inertia, *I*_*xx*_ and *I*_*zz*_, respectively, (**g**) similar to those in birds. **h**,**i**, Location of the CG (**h**) and moments of inertia (**i**) are quite constant with the tail shape morphing and twist. **j**, Wing and tail shape morphing controls longitudinal stability by altering location of the neutral point. **k**,**l**, Lift and drag (**k**), and pitching moment (**l**) generated by wing and tail shape morphing. Solid lines and shaded areas indicate mean value and standard deviation, respectively.

In contrast to fixed-wing aircraft’s tail, which is used solely for stability and control and is positioned sufficiently away from the wings to minimize negative effects of wing-induced wakes^19^, avian tails are located comparatively closer to the wing trailing edges and they play important aerodynamic functions, such as lift augmentation^10,20^, drag reduction^21–23^, and pitch control^24^. Additionally, birds compensate for the absence of vertical tail stabilizers found in aircraft by twisting their tails around the body axis^2,5,6,25,26^. Similar to birds, the morphing tail used here is located close to the drone wings, does not have a vertical rudder, can independently twist around the body axis (Fig. 1e), and can also change pitch angle for longitudinal control^13,27,28^ (Extended Data Fig. 2). As the size and shape of the tail vary substantially across bird species^29^, we designed the tail used in this study with the length and shape similar to those of soaring hawks of comparable scales^30^ (‘Feathered wing and tail design’ in Methods and Extended Data Fig. 2).

In birds, sweep morphing of aerial surfaces alters not only aerodynamic but also inertial characteristics, two quantities that substantially impact flight manoeuvrability^15^. Similarly to raptors during gliding with fully extended wings, the drone’s Center of Gravity (CG) lies behind the wing aerodynamic center, which is located approximately at the quarter of the wing mean chord^15,31^. When morphing, the CG shifted only by 3% of the full body length (Fig. 1f), similarly to birds^15^. Whereas, wing extension increased roll and yaw moments of inertia by up to 3-fold and 1.8-fold, respectively (Fig. 1g), consistently with observations of cooper’s hawks and peregrine falcons^15^ during the elbow and wrist extension (approximately 4-fold and 2-fold, respectively). Instead, tail sweep morphing and twisting did not significantly affect moments of inertia due to the comparatively low mass of the tail, which represents less than 3% of the body mass (Fig. 1h,i).

We then investigated static aerodynamic performance of the drone in wind tunnel experiments (Fig. 1j-l, ‘Aerodynamic characterizations’ in Methods, and Extended Data Fig. 4). Wing morphing altered lift and drag generation, and affected pitch stability (Fig. 1j-l), similarly to birds^15^. By fully extending the wings with a fully folded tail, the neutral point lied ahead of the CG, causing the drone to be slightly unstable (*C*_*m,α*_ *= ∂C*_*m*_*/∂α* > 0, where *C*_*m*_ is the pitch coefficient and *α* is the angle of attack). Folding the wings shifted the neutral point aft of the CG, thus shifting the drone to a stable longitudinal mode (*C*_*m,α*_ < 0), but decreased lift. In addition, we found that spreading the tail also enabled the drone with fully extended wings to transition from unstable to stable states, while increasing lift (Fig. 1k,l). These results suggest that birds may prefer to use tail spread morphing rather than wing sweep morphing to control longitudinal stability when flying at low speed where lift is a priority. In summary, these results indicate that the avian-informed drone displays inertial and aerodynamic properties similar to those of birds, and represents a suitable model for studying avian turning manoeuvres.

### Tail twist controls steady banked turns

We started investigating the role of tail twist in banked turns by conducting aerodynamic characterization in an open wind tunnel (Fig. 2, ‘Aerodynamic characterizations’ in Methods). We found that tail twist contributed to all 6-axis forces (lift, drag, and side force) and moments (pitch, roll and yaw moments) of the drone. Twisting the tail around the body axis (regardless of the rotation direction) reduced its longitudinal surface, thus reducing lift (Fig. 2a). The reduced tail lift caused the drone to experience a nose-up orientation and shift from longitudinally stable to unstable mode (Fig. 2b). During a banked turn, a traditional aircraft rolls around its body axis to redirect the lift vector into the direction of the roll that adds horizontal component providing centripetal acceleration to execute the turn, but decreases the vertical lift component and thus flight altitude. To gain more lift, most aircraft deflect the elevator at the tail to generate a pitch-up moment, which increases the wing angle of attack^32^. Thus, the nose-up orientation induced by the tail twist of the avian-informed drone may serve a similar role by increasing the overall lift force during the turn.

**Fig. 2.**
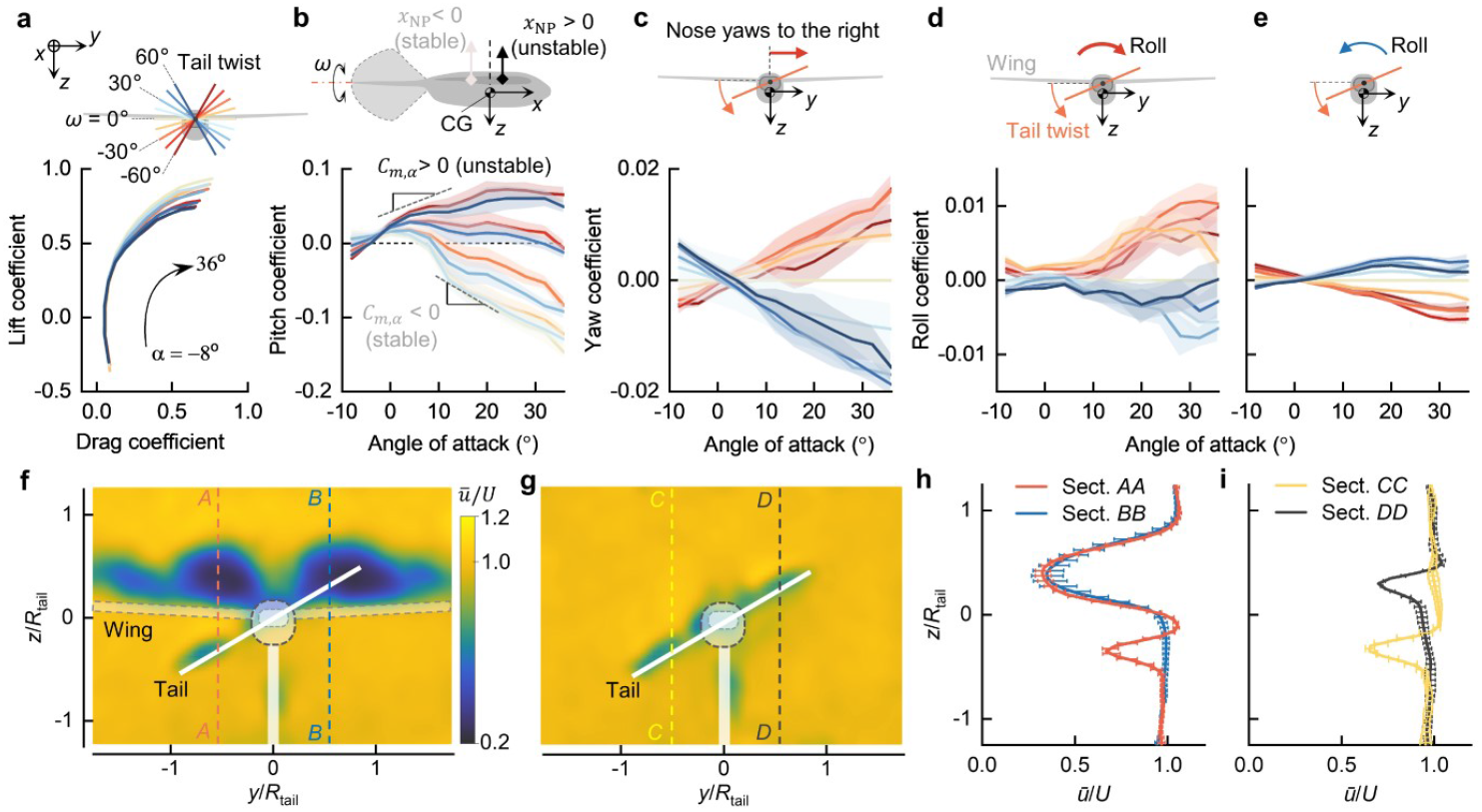
Aerodynamic characteristics of the feathered drone with tail twist. **a**, Twisting the tail reduces the longitudinal surface area, thus reducing lift. **b**, Pitching moment generated as a result of tail twist. Reduced lift from the tail shifts the drone from stable to unstable configurations by generating more pitch-up moment. **c**, Tail twist generates a proverse yaw moment that supports the turn. **d**,**e**, Roll moments with presence (**d**) and absence (**e**) of the wings. **f–i**, Twisting the tail sends the upper tail part into the turbulent wake of the wing. **f**,**g**, Velocity maps at 50% chord of the tail with presence (**f**) and absence (**g**) of the wings. **h**,**i**, Velocity profiles at y = 0.5*R*_tail_ = 0.11 m with presence of the wings shown in **f** at the sections *AA* and *BB* (**h**) and absence of the wings shown in **g** at the sections *CC* and *DD* (**i**). The measurement was conducted at the inflow velocity of *U*_ref_ = 7 m/s with the angle of attack of *α* = 16°. In **a–e**, solid lines and shaded areas indicate mean value and standard deviation, respectively.

Twisting the tail tilted the tail lift vector and therefore generated side forces, resulting in yaw moments, namely a negative tail twist produced a positive yaw moment (nose yaws to the right), and vice versa (Fig. 2c). We also found that tail twist generated roll moments, namely negative twist resulted in a positive roll moment (right wing downwards), and vice versa (Fig. 2d), consistently with observations of birds^1,3^. Taken together, these data show that tail twisting enables roll control required to initiate the banked turn and generates yaw moment that counters the adverse yaw induced by the higher drag of the outer wing during turning. However, when the wings were removed, twisting the tail generated roll moments in the same direction of the tail motion (Fig. 2e). To gain insight into how wings affect the roll moment generated by the tail, we conducted flow visualization experiments (Fig. 2f-i, and ‘Flow visualization’ in Methods). We found that, when twisted, the upper side of the tail entered the low-speed wake region induced by the wings, thereby reducing its force generation (Fig. 2f,h); in contrast, the other side of the tail, which descended away from the wing, was not affected by the wing-induce wake. Consequently, asymmetric lift between the two sides of the tail generated a roll moment. On the other hand, without the wing ahead, there was no significant difference in the flow fields between the two sides of the tail (Fig. 2g,i). As the CG location was lower than the pivot point of the tail (rotation axis) (Fig. 2e), twisting the tail increased the moment arm to the upper half and reduced that in the lower half of the tail. As a result, roll moment was generated in the same direction of the tail twist. We also confirmed that this roll moment was negligible when the CG was located at the tail rotation axis (Extended Data Fig. 5).

To test whether the moments generated by tail twist are sufficient to initiate banked turns, we conducted indoor flight experiments (Fig. 3, ‘Flight experiments’ in Methods, Extended Data Figs. 6 and 7, and Supplementary Videos 1-3). We replicated the banked turn strategy observed in birds by a two-phase tail twist^3^ (Fig. 3a,b). The results showed that the drone could perform banked turns with a turning radius (=1/curvature) of 10.8 ± 3.0 m at a bank angle of 31.2 ± 8.5° (Fig. 3d-l) with roll and yaw motions (Fig. 3g,i), and nose-up pitch (Fig. 3h) induced by tail twisting (Fig. 3b). To complete the turn, the drone temporarily reversed its tail twist and returned to level flight; if no action was taken, the drone kept increasing its roll and yaw while decreasing pitch (Fig. 3g-l) and as a result, the drone tumbled (Supplementary Video 3). We also found that the reaction force induced by the tail twist acceleration did not induce a rotation of the drone in the opposite direction (Extended Data Fig. 8). These results suggest that birds may stabilize the exit phase of banked turns by temporarily reversing the tail twist direction instead of relying only on inherent lateral stability^1^.

**Fig. 3.**
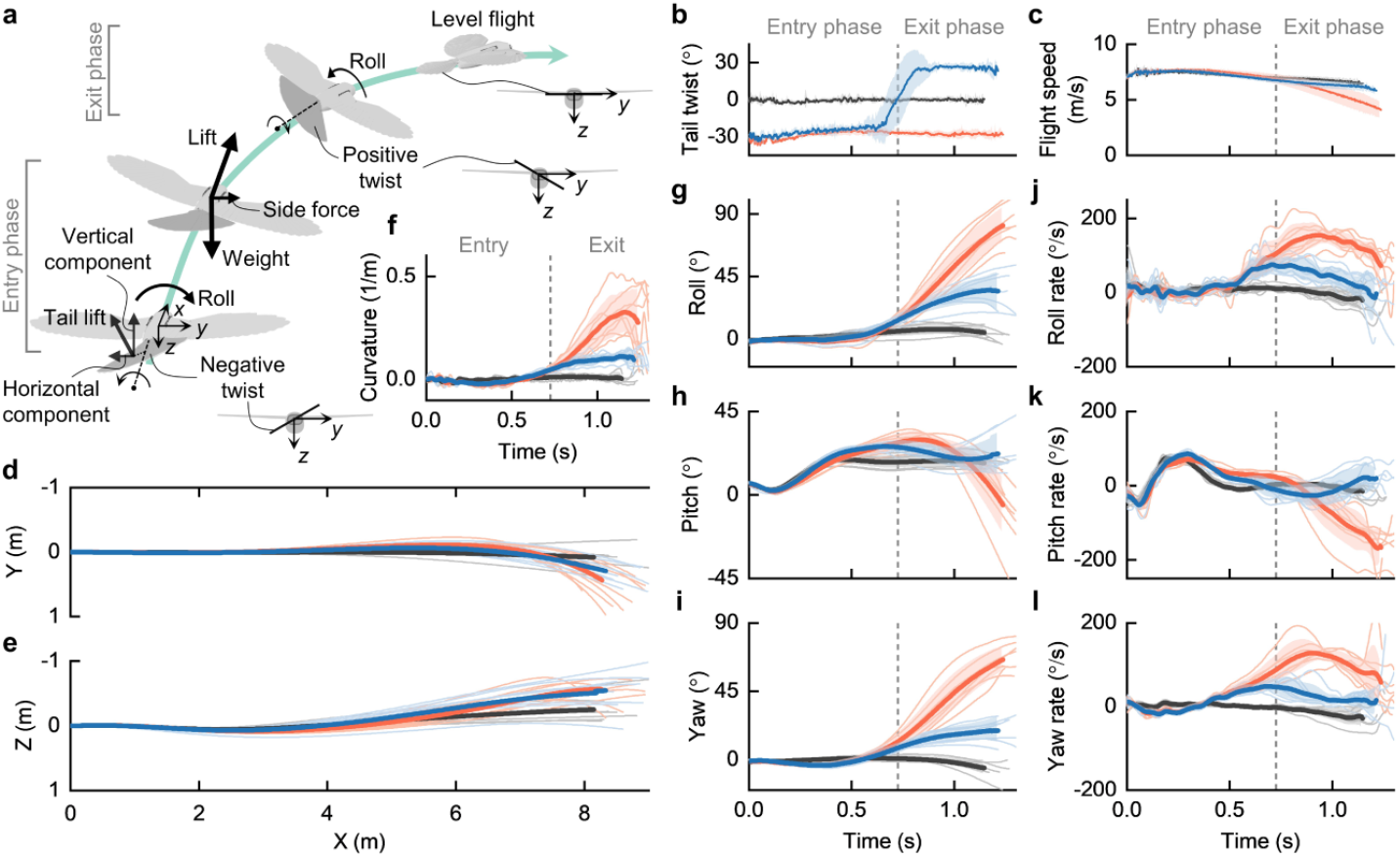
Flight experiments confirm that drones and birds can merely twist their tails to execute steady banked turns. **a**, Two-phase turning manoeuvres. Negative tail twist initiates a right turn and reverses to positive twist to exit the turn. **b**, Tail-twist angles in three flight patterns: trimmed flight (n = 7, black), banked turns without (n = 8, red) and with (n =8, blue) reversing the tail twist at the end of the turn. Tail twist angle is set to be ± 30°, which generates maximum rolling moments shown in Fig. 2d. **c**, Flight speeds during turning. **d**,**e**, Flight trajectory in the *XY*-plane (**d**) and the *XZ*-plane (**e**). **f**, Curvature of the turns. **g**–**i**, Body attitude angles: roll (**g**), pitch (**h**), and yaw (**i**). **j**–**l**, Body rates: roll rate (**j**), pitch rate (**k**), and yaw rate (**l**). In **b** to **l**, thin lines represent individual flight trials; thick lines and shaded areas are the mean and standard deviation from the mean, respectively.

### Tail twist controls high-speed sharp turns

Flying at high speed through cluttered environments requires sharp turns to avoid obstacles, but the role of tail twist in such turning manoeuvres is still unexplored. Although birds can resort to either flapping or gliding manoeuvrings in a turn^33^, flapping is observed in birds flying at low speed^34–36^ and expends much more metabolic energy than gliding manoeuvring^37,38^. Whether and how birds execute high-speed sharp turns by gliding manoeuvring remains unclear. As the turning radius is proportional to the square of the flight speed^32,39^, performing a high-speed turn with small turning radius may require quick deceleration before the turn. Many large birds engaged in agile manoeuvres, such as perching^40^ and recovery from high-speed dive^41,42^, decelerate by executing a rapid pitch-up manoeuvre to transfer kinetic energy to potential energy. Here we hypothesize that birds may leverage a similar strategy, and coordinate tail pitching and twisting to perform sharp manoeuvres when flying at high speeds.

We conducted flight experiments on high-speed sharp turning by applying a coordination of rapid pitch-up manoeuvre and tail twist (Fig. 4, ‘Flight experiments’ in Methods). To initiate the pitch-up manoeuvre, the tail was deflected upward^9,40,43^, causing the drone to climb (Fig. 4a,e). During the climb, negative tail twist was applied to induce roll and yaw movements to the right. When the climbing phase reached peak altitude, the drone reversed the tail twist to exit the turn. We found that the coordination allowed the drone to perform sharp 90° turns with small turning radii. Experimental results showed that the drone flying at an initial speed of 8.8 ± 0.5 m/s could turn with a radius of 1.9 ± 0.3 m and a bank angle of 84.4 ± 5.2° (Fig. 4b-m (black), and Supplementary Videos 4). The pitch-up response allowed the drone to reduce its speed to 2.9 ± 0.3 m/s at peak altitude (Fig. 4c), thus converting 24.2 ± 2.7% kinetic energy into potential energy, while dissipating 64.7 ± 2.3% through aerodynamic braking (Fig. 4g). Nevertheless, tail twist alone could slowly recover the roll motion to level flight in the exit phase of the turn (Fig. 4h,k, black). To address this issue, we added asymmetric wing folding observed in turning birds^1,3^ (Fig. 4a, and Extended Data Fig. 9) and obtained a faster level flight recovery after the turn (Fig. 4h,k, and Supplementary Videos 5). However, the faster recovery came at a cost of increased turning radius of 3.2 ± 0.6 m. By adjusting the starting time of the asymmetric wing folding during the exit phase, it is possible to regulate the turning radius and heading direction (Extended Data Fig. 10). However, the drone could not turn with asymmetric wing folding without tail twisting due to adverse yaw effect (Extended Data Fig. 11, and Supplementary Videos 6). These results indicate that tail twisting could play a major role also in sharp turning manoeuvres at high flight speed.

**Fig. 4.**
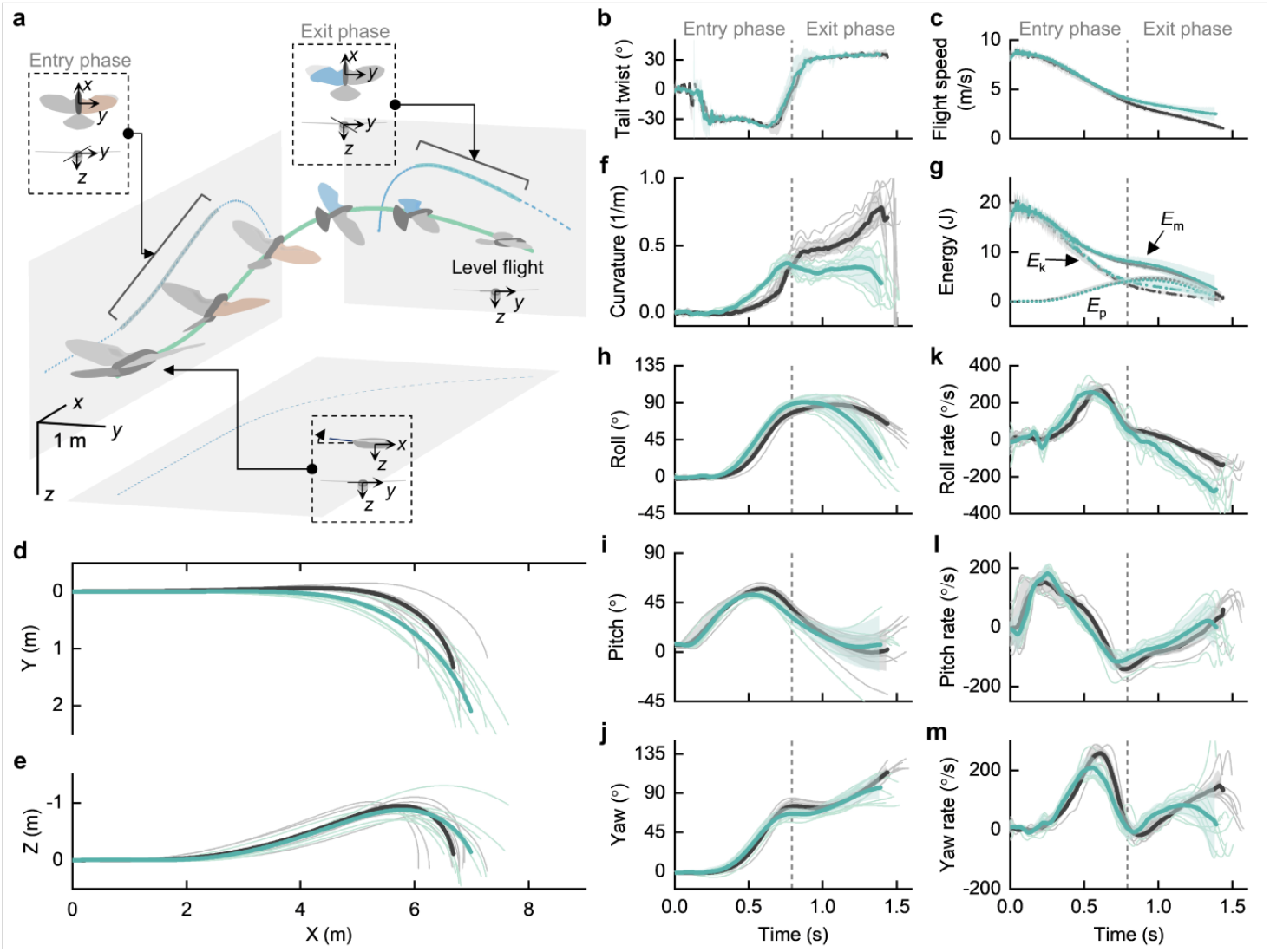
High-speed sharp turning manoeuvres enabled by tail twist with (green) and without (black) coordination of wing shape morphing. **a**, Illustration of the control input sequence to perform a sharp turn with the tail-twist and wing morphing coordination. For the right turn, the drone deflected the elevator upward to execute a rapid pitch-up manoeuvre. Thereafter, negative tail twist and right wing folding were synchronously applied to create roll and yaw motion. At the peak of the altitude, positive tail twist and left wing folding were applied to return to the level flight. **b**,**c**, Tail twist angles (**b**) and flight speeds (**c**) in flights with (n =10, green) and without (n =8, black) coordination of wing shape morphing. **d**,**e**, Flight trajectories in the *XY*-plane (**d**) and the *XZ*-plane (**e**). **f**, Curvature of the turns. **g**, Kinetic (*E*_k_), potential (*E*_p_), and mechanical (*E*_m_ = *E*_k_ + *E*_p_) energies during the turn. **h**–**j**, Body attitude angles: roll (**h**), pitch (**i**), and yaw (**j**). **k**–**m**, Body rates: roll rate (**k**), pitch rate (**l**), and yaw rate (**m**). In **b** to **m**, thin lines represent individual flight trials; thick lines and shaded areas are the mean and standard deviation from the mean, respectively.

## Conclusions

These results suggest that tail twisting can play a substantial role in turning manoeuvres at both low and high speeds. When twisted, the tail enters asymmetric flow regions induced by the wings, resulting in the generation of both roll and yaw moments that initiate, regulate, and end a turn. Simultaneously, reduced lift at the tail induces a nose-up motion that increases the wing angle of attack, thus generating more lift to compensate for the loss induced by the banking motion and thus help preserve the flight altitude. We propose that, among other control methods^4,13,32,39,44^, tail twisting may be the most efficient approach for controlling steady turning manoeuvres in birds, as it requires activating only one group of tail muscles. Other methods that rely on asymmetric wing morphing or wing twisting may generate more effective roll moment (Extended Data Fig. 9), but necessitate additional muscular controls to counteract adverse yaw^4,32,44^.

Furthermore, we showed that sharp turns at high speed can leverage tail twisting in combination with a rapid pitch-up manoeuvre that not only substantially reduces flight speed, but also enhances the effectiveness of roll moments generated by the tail twist (Fig. 2d). The results also showed that faster return to level flight after a turn can be achieved by coordinating tail twisting with asymmetric wing morphing, although the low flight speed at this stage may force birds to use powerful flaps to regain initial flight conditions.

Taken together, our findings with an avian-informed drone contribute to a better understanding of avian turning behaviours that are difficult to study with animals in controlled laboratory experiments. Although there are gaps in the morphological complexity and range of motion when compared to birds, the aerodynamic forces generated by the feathered drone are comparable to those of a hawk wing^45,46^ (Extended Data Fig. 4). Finally, the results described here have implications for effective control of drones with morphing aerial surfaces, thus broadening their range of applications across diverse environments, from open spaces to complex confined areas such as cities or dense forests.

## Methods

### Feathered wing and tail design

#### Wing skeletal architecture

The mechanical wing skeletal structure was designed adapting morphological parameters of a hawk wing skeleton. It is composed of three segments (humerus-, ulna-, and manus-like linkages) made of 1 mm glass fiber composite sheet that can rotate around the shoulder, elbow, and wrist joints (Extended data Fig. 1). The motion of the three segments is coupled through two pulley-string mechanisms, enabling the wing to be actuated by only one servomotor. The first pulley, with its radius *r*_1_, is fixed to the body frame at the shoulder joint and connected to the second pulley with radius *r*_2_, which is fixed to the ulna-like linkage at the elbow joint, through string 1. Simultaneously, the third pulley with radius *r*_3_ fixed to the humerus-like linkage and rotated around the elbow joint, is connected by string 2 to the fourth pulley with radius *r*_4_. This last pulley is affixed to the manus-like linkage and rotated around the wrist joint. Thus, range of motion of each segment can be determined by

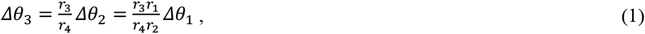

where *θ*_1_, *θ*_2_, and *θ*_3_ are the sweeping angles of the humerus, ulna, and manus linkages, respectively (Extended data Fig. 1), and 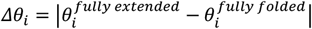, where *i* = 1,2,3.

#### Wing feathers

The wing is formed by 20 lightweight artificial feathers, including 9 primary (P1 – P9) and 11 secondary (S1 – S11) feathers, as shown in Extended data Fig. 1e. Inspired by a bird feather, the artificial feather is composed of a 1-mm S-shape vane made of flexible expanded polypropylene (EPP) foam with a density of 45 kg/m^3^ and a reinforced calamus or quill made of a 1.5 mm carbon fiber tube. All the feathers, except for the primary feather P9 that is fixed to the tip of the manus-like linkage, are attached to feather joints on the skeletal structure that could passively rotate in the wing plane (Extended data Fig. 1). The carbon shafts of the feathers are interconnected by a non-elastic tendon with one end attached to the body frame and the other end fixed to the manus-like linkage. Through this design, the 20 feathers formed a configuration vector **q** with dim[**q**] = 20 are underactuated via wing skeletal control with a single degree of freedom^44,47^. The feathers’ ranges of motion are also constrained to ensure that all the feathers still overlap consistently to form a complete surface when the wing is fully extended (Extended data Fig. 1g).

#### Wing covert

Birds overlay their wing skeletons by numerous overlapping covert feathers to streamline the wing. Replicating such coverts may cause mechanical complexity and increased wing mass. To replace the role of the coverts, we covered the mechanical wing skeletal structures and the feather shafts with discrete lightweight layers made of EPP foam (20 kg/m^3^). The three covert-inspired layers overlap each other at the elbow and wrist joints to facilitate morphing (Extended data Fig. 1e).

#### Tail structure and feathers

The tail is capable of controlling independently four degrees of freedom including shape morphing, twisting, elevator deflection, and lateral deflection, which was deactivated in this study (Extended data Fig. 2). It is formed by 11 overlapping feathers that distribute symmetrically across the body’s symmetric plane (*xz*-plane). Except for the fixed central feather with symmetric anterior and posterior vanes, the other tail feathers, which have a similar shape to the wing feathers, can rotate on the tail plane around feather joints located at the tail skeletal structure made of 0.5-mm glass fiber composite sheet. The feather shafts are interconnected by a pre-stretched 3D-printed elastic tendon (NinjaFlex 85A). We used a servomotor to actively and synchronously actuate the two outermost feathers through a non-elastic fishing cable (diameter of 0.4 mm, Spiderwire Dura-4 Braid, 45 kg) running through the center of the universal tail joint and along the inside of the hollow rectangular body shaft^28^. The tail spread is thus activated by pulling the cable using the servomotor, whereas the elastic force of the tendon triggers the tail fold. Similar to the wing, the morphing tail is an underactuated system. The maximum tail spread angle is 120°, which was estimated to provide optimal lift-to-drag ratio^29^.

The tail can rotate around a universal joint that enables both elevator and lateral deflection controls^28^, actuated by two servomotors affixed to the body shaft. To control tail twist, we split the body shaft into anterior and posterior parts. Whilst the anterior part is fixed to the fuselage, the posterior part―where the tail is located―can rotate around it via a bearing (Extended data Fig. 2). By connecting this rotating shaft to a servomotor through a pulley-string mechanism, tail twist is fully actuated with a maximum twisting angle of ±60°.

### Drone fabrication and avionics

We fabricated glass fiber composite sheets using a CO_2_ laser cutter (Trotec Speedy 400) for the wing and tail skeletal structures, as well as the supported frames that link the wing bases to the body shaft made of 6 mm × 6 mm carbon fiber square tube (Extended data Fig. 3). All the pulleys, joints, and servomotor holders were 3D printed in acrylonitrile butadiene styrene (ABS) using Stratasys Dimension Elite printer. The fabricated parts were then assembled manually and bonded together by cyanoacrylate super glue. We connected the pulleys located at the wing joints using 0.4-mm durable strings (Spiderwire Dura-4 Braid, 45 kg) to create the pulley-string mechanisms that drive the wing shape morphing. We fabricated the EPP foams for the feather vanes, coverts, and body cover using a hot wire machine (CNC-Multitool CUT 1620S). The feather vane was attached to the 1.5-mm carbon fiber quill using the UHU Por glue. The quill was then inserted into the feather joint, which can rotate around a 1.5-mm carbon shaft in the skeletal structure.

To actuate the wing and tail morphing, we used KST X08plus V2 servomotors (KST digital servo, 5.3kg.cm at 8.4V). For propulsion, we used a brushless DC motor (T-Motor AT-2306 KV2300), a 8 × 4.5 inch GWS propeller, a Pulsar A-15 speed controller, and a 3-cell 900 mAh lithium polymer battery. An 8-channel FrSKY RX8R receiver was used to communicate with the pilot through the Taranis X9D Plus transmitter. The mass of the drone with a wingspan of 1.1 m and full body length of 0.54 m is 460 g.

### Aerodynamic characterizations

#### Wind tunnel experiments

We characterized aerodynamics of the feathered drone in a 2 × 1.75 m open wind tunnel (WindShape, turbulence intensity < 1%) made of an 8x7 array of controllable fan modules at the mean wind speed of 7 m/s, which is similar to the gliding speeds of hawks^24,27^. To acquire the aerodynamic forces and torques, we placed the drone on a calibrated 6-axis force/torque sensor (ATI Nano 25, force resolution 0.0625 N, torque resolution 0.75 mNm) mounted at the extended tip of a robotic arm (TX-90, Stäubli) positioned in front of the wind tunnel. The robotic arm is programmable to systematically change the angle of attack of the drone from -8° to 36° at 4° intervals. The force/torque sensor was located at the CG of the drone. The force/torque signals were logged by the NI-DAQmx 9.5.1 logger (National Instruments) through the installed ATI DAQ F/T software (ATI Industrial Automation) at the sampling rate of 3000 Hz with an averaging level of 5. At each angle of attack, we recorded the data for 8 s (4800 data points). Before starting the measurement, we zeroed all forces and torques at a 0° angle of attack and no-speed condition. For constant power input, we used an external DC power supply (PeakTech 6226) to power the servomotors for controlling different wing and tail morphing configurations. The propeller was removed during the measurement; therefore, the results exclude effects of propeller slipstream, considering similar flight conditions as in birds. We first validated lift and drag generation of the feathered drone by comparing them to those of a hawk wing within a similar range of Reynolds number^45^. We then measured the forces and torques generated by various wing shape morphing both symmetrically and asymmetrically, tail shape morphing, and tail twisting.

We also conducted wind tunnel experiments to investigate the effect of rate of change in tail twist on the roll motion (Extended data Fig. 8). We placed the drone on the extended tip of the robotic arm, which allows only one degree of freedom in roll. We then placed markers on the drone and used a motion tracking system (26 Optitrack cameras, 240 Hz) to monitor its roll motion in response to the tail twist inputs. The drone was commanded to twist its tail from 0 to -30° with step, 0.5-s, and 1.0-s slowdown inputs.

#### Data processing

We analyzed the force and torque data over 4-s period (2400 data points) at each angle of attack, excluding those at the transition stage between them. The force/torque sensor can record three force (*F*_x_, *F*_y_, and *F*_z_) and three torque (roll: *M*_x_, pitch: *M*_y_, and yaw: *M*_z_) components in the *xyz* body coordinate system. By using the measured *F*_x_ and *F*_y_, we calculated lift (*L*) and drag (*D*):

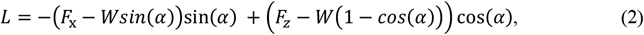

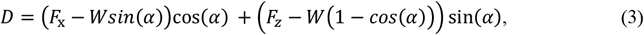

where *W* is the weight of the drone used in the measurement, and *α* denotes the angle of attack. Non-dimensional lift coefficient (*C*_*L*_) and drag coefficient (*C*_*D*_) were then calculated as

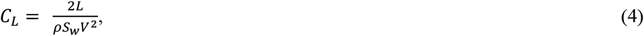

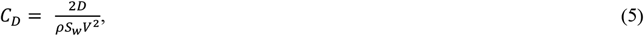

where *ρ* denotes the air density, *S*_w_ is the wing area of the fully extended configuration, and *V* is the input flow speed. Thereafter, we obtained pitch, roll and yaw torques in coefficient form:

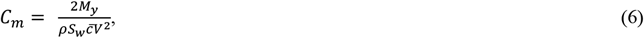

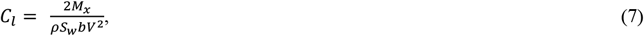

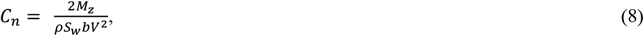

where 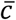 is the mean wing chord and *b* is the tip-to-tip wingspan.

### Flow visualization

We used a Probe Capture (ProCap) with a hand-guided 5-hole pressure probe for real-time visualization and measurement of the airflow field^48^. We fixed the drone on the extended tip of the robotic arm at an angle of attack of 16°, positioned in front of the wind tunnel with an inflow speed of 7 m/s. We defined a visualization plane (0.7 m × 0.5 m) perpendicular to the airflow direction and located at the middle chord of the tail. The instantaneous position of the probe, tracked by a motion tracking system (26 Optitrack cameras) at a tracking frequency of 240 Hz, was synchronized with the inflow data streamed from the probe. We conducted measurements for the fully extended tail at a twisting angle of -30° with and without the presence of the wings. The measured data were post-processed in an open source multi-platform software (ParaView 5.11.1) for data analysis and visualization.

### Flight experiments

Flight experiments were conducted in a 10 m × 10 m × 8 m indoor flight arena equipped with a safety net and a motion tracking system (26 Optitrack cameras, 240 Hz) that can track the reflective markers located on the drone (Extended data Fig. 6). We launched the drone using a custom-built, programmable linear launcher at speeds of approximately 7 m/s for steady turns and 9 m/s for sharp aerobatic turns. We performed experiments on both the feathered and fixed-wing drones with similar morphologies that are capable of tail twisting. We defined a fixed *XYZ* coordinate system located at the tip of the launcher, in which the *XY*-plane is the horizontal plane, and the *Z*-axis points downward. We first trimmed the drones to maintain a straight flight path and similar body attitudes without any control inputs (Fig. 3b-m; black). We set the throttle at 50% to maintain a consistent flight speed of 7 m/s of the drone after released from the launcher.

To test the open-loop response of the drone to tail twist under a steady turn scenario, we automated flight trials by predetermining tail-twist control inputs while locking other control surfaces. For the case of single-phase tail twist (Fig. 3b-m; red), the tail was locked at a twist angle of -30° throughout the entire flight. In the case of two-phase tail twist (Fig. 3b-m; blue), the flights were initiated with the tail-twist angle set at -30° until the drone reached a banked angle of approximately 20°. Thereafter, the tail was switched to the twist angle of 30° to recover the level flight.

For sharp aerobatic right turns, we launched the drone with an upward tail deflection of -10° to execute the pitch-up manoeuvre after released from the launcher. When the pitch angle reached about 20° (we assumed that the pitch angle was similar to the angle of attack at this initial stage), a negative tail twist of -30° was applied to roll and yaw the drone to the right. Note that, to ensure a desired flight turn, the tail twist should be activated after the drone reaches a pitch angle that is larger than the magnitude of the tail deflection angle. Otherwise, adverse yaw may be occurred. At the peak of the flight altitude when the drone achieved desired roll and yaw angles, we reversed the tail twist to 30° to return to the level flight. All these processes were automated through predetermined control inputs.

From the measured flight data extracted by the motion tracking system, we calculated the curvature (*κ*) and radius (*r*) of the horizontal turn^49^:

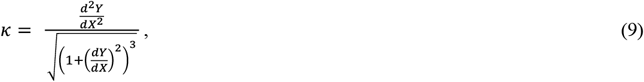

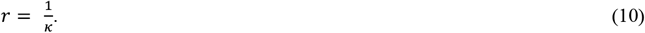

Finally, we obtained mechanical (*E*_m_), kinetic (*E*_k_), and potential (*E*_p_) energies from the following equation

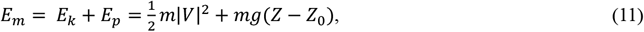

where 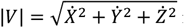 is the flight speed (m/s), and *m* is the mass of the drone (kg).

## Acknowledgements

This project was partially funded by the European Union’s Horizon 2020 research and innovation programme under grant agreement ID: 871479 AERIAL-CORE, and by Armasuisse grant nr. 8003529756 “Wind Riders”. We thank E. Ajanic for comments on the drone design and his previous work on LisHawk drone, and V. Wüest, S. Jeger, M. Askari and R. Zufferey for designing and building the drone launcher.

## Author contributions

H.-V.P. conceived the idea and designed the research, built the robot, performed experiments, processed and analyzed the data, and wrote the manuscript. D.F initiated the project, and contributed to the research design, data analysis and writing of the manuscript. Both authors gave final approval for publication.

## Competing interests

The authors declare no competing interests.

---

**Extended Data Fig. 1.**
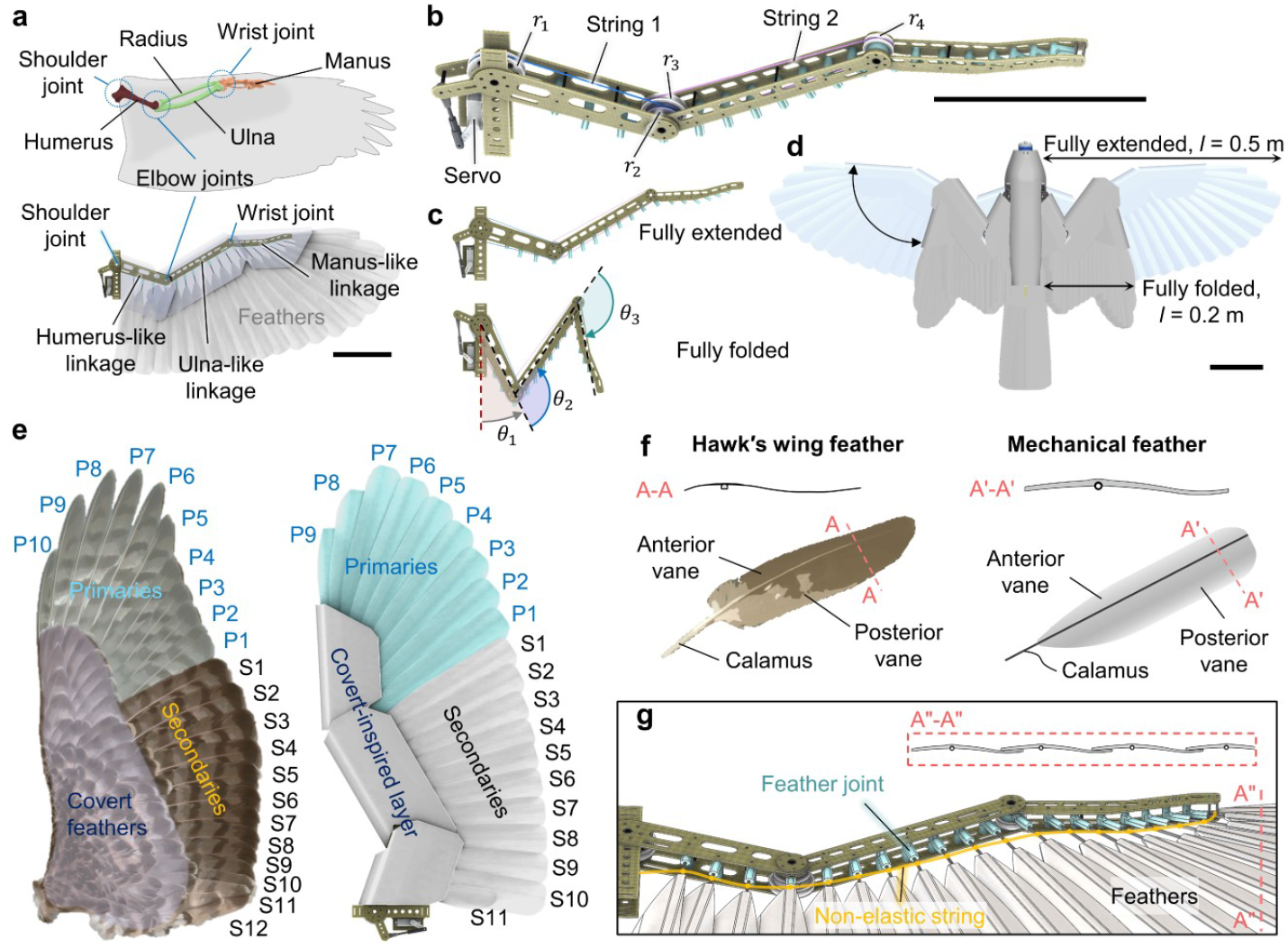
Bio-informed wing skeletal and feather design. **a**, Morphing wing design with multi-segment, multi-joint skeleton inspired by bird wing (top). **b**, Detail of the wing skeleton driven by a single servomotor through two coupled pulley-string mechanisms. **c**, Wing skeleton in fully extended and fully folded configurations. **d**, The bird-inspired morphing wing enables the drone to change wingspan and surface area. **e**, The wing is formed by primary-, secondary- and covert-inspired feathers, resembling those of a bird wing. Image of bird wing (Northern Goshawk, *Accipiter gentilis*) courtesy of Puget Sound Wing and Tail Image Collection (Accessed on 04/12/2023). **f**, Feather design with a bird-inspired vane and calamus or quill. **g**, The feathers are connected to the skeletal structure through in-plane rotating joints. The quills of the feathers are interconnected though a non-elastic tendon (yellow) to ensure that all the feathers still overlap consistently to form a complete surface during wing morphing.

**Extended Data Fig. 2.**
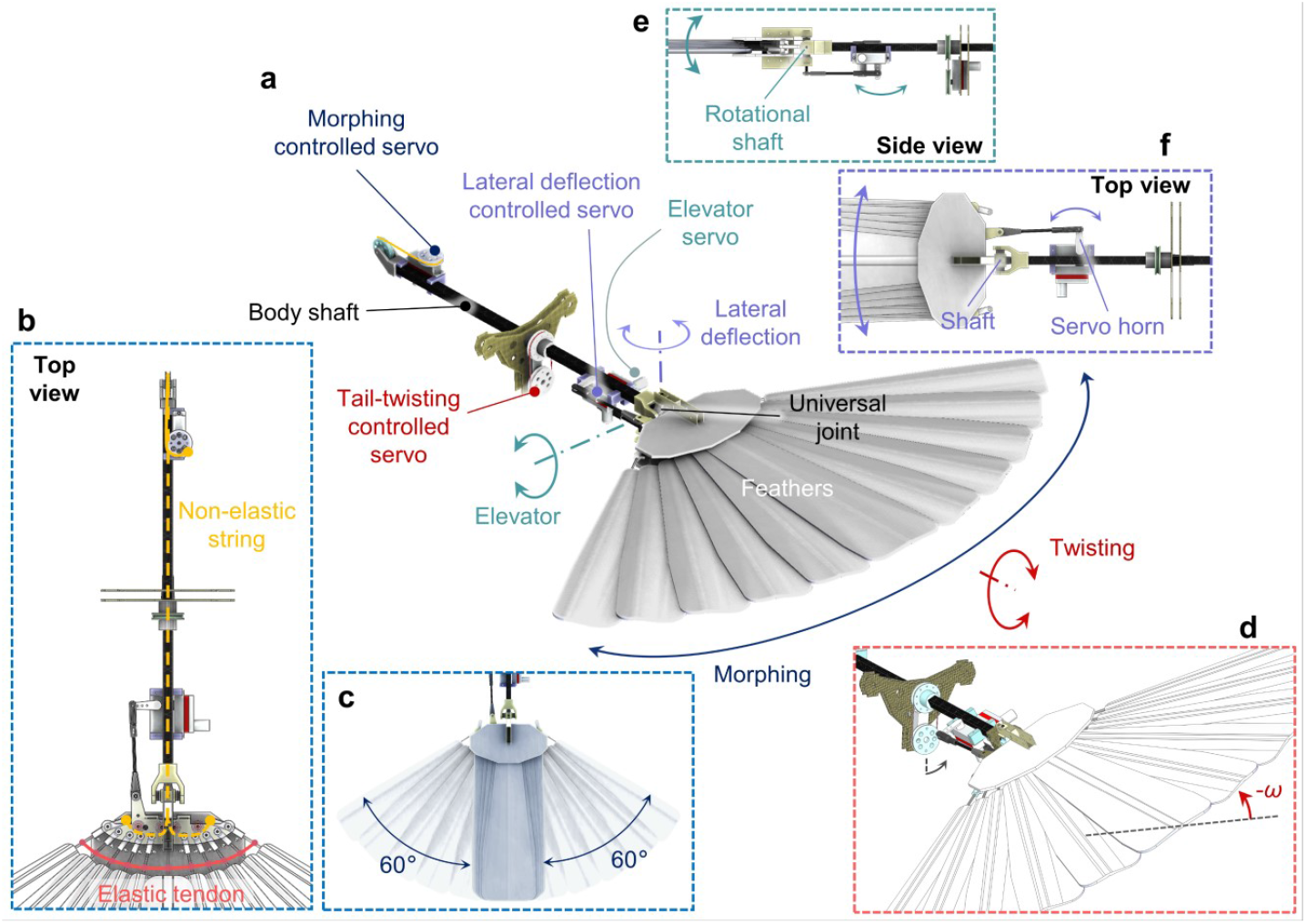
Rudderless morphing tail design. **a**, The tail formed by 11 overlapping feathers is capable of shape morphing, twisting, elevator deflection, and lateral deflection. **b**, Top view of the tail shape morphing mechanism. **c**, The tail in fully folded and fully spread configurations. **d**, Mechanism of tail twisting. **e**, Side view of the elevator-controlled mechanism. **f**, Top view of the lateral deflection-controlled mechanism. This tail motion was locked in this study.

**Extended Data Fig. 3.**
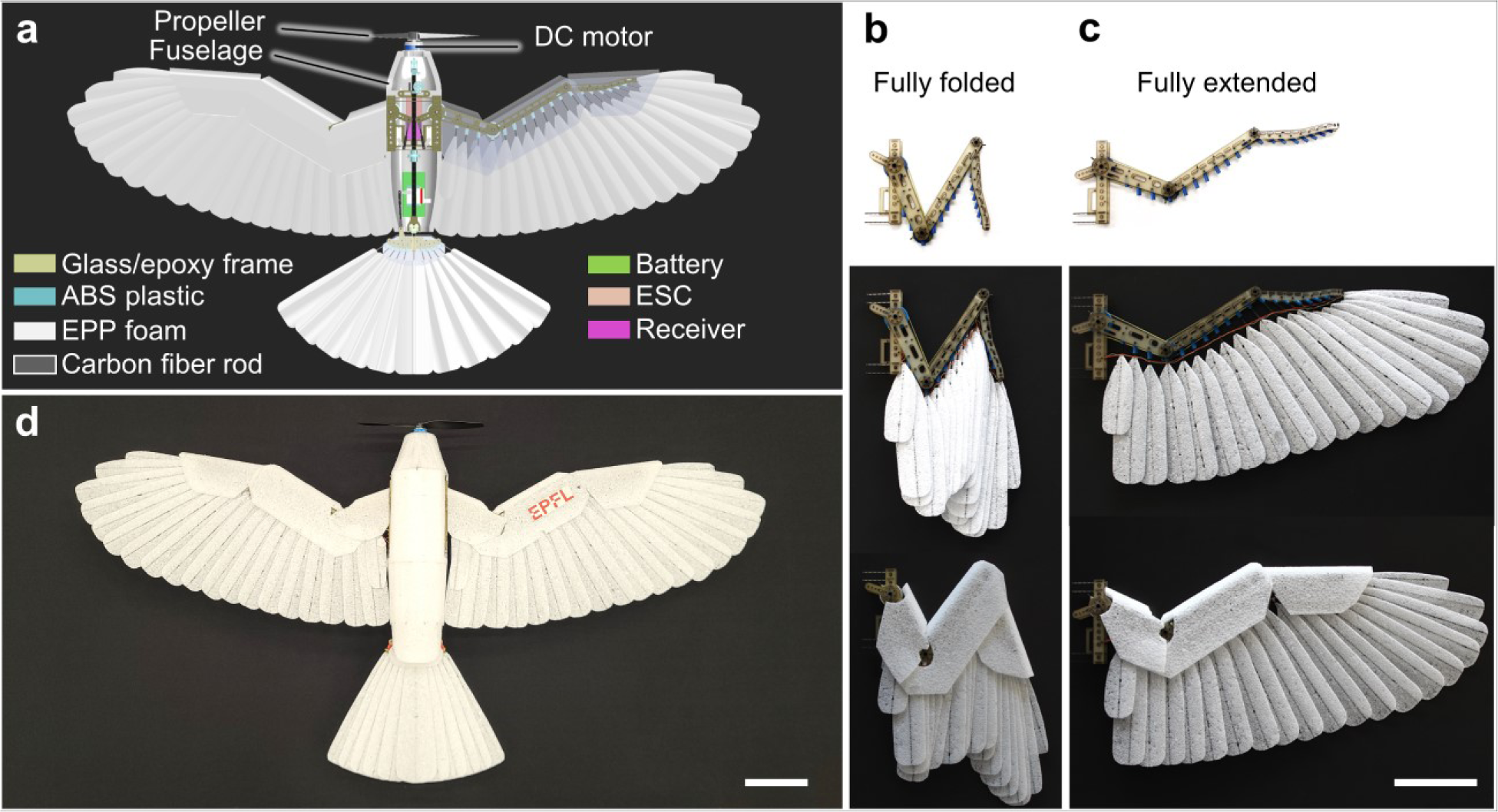
Drone prototype. **a**, Overview of materials and electronics used to fabricate the drone. **b**,**c**, Fabricated wing in fully folded (**b**) and extended (**c**) configurations. **d**, The drone prototype with fully extended wings and half-extended tail. Scale bar: 10 cm.

**Extended Data Fig. 4.**
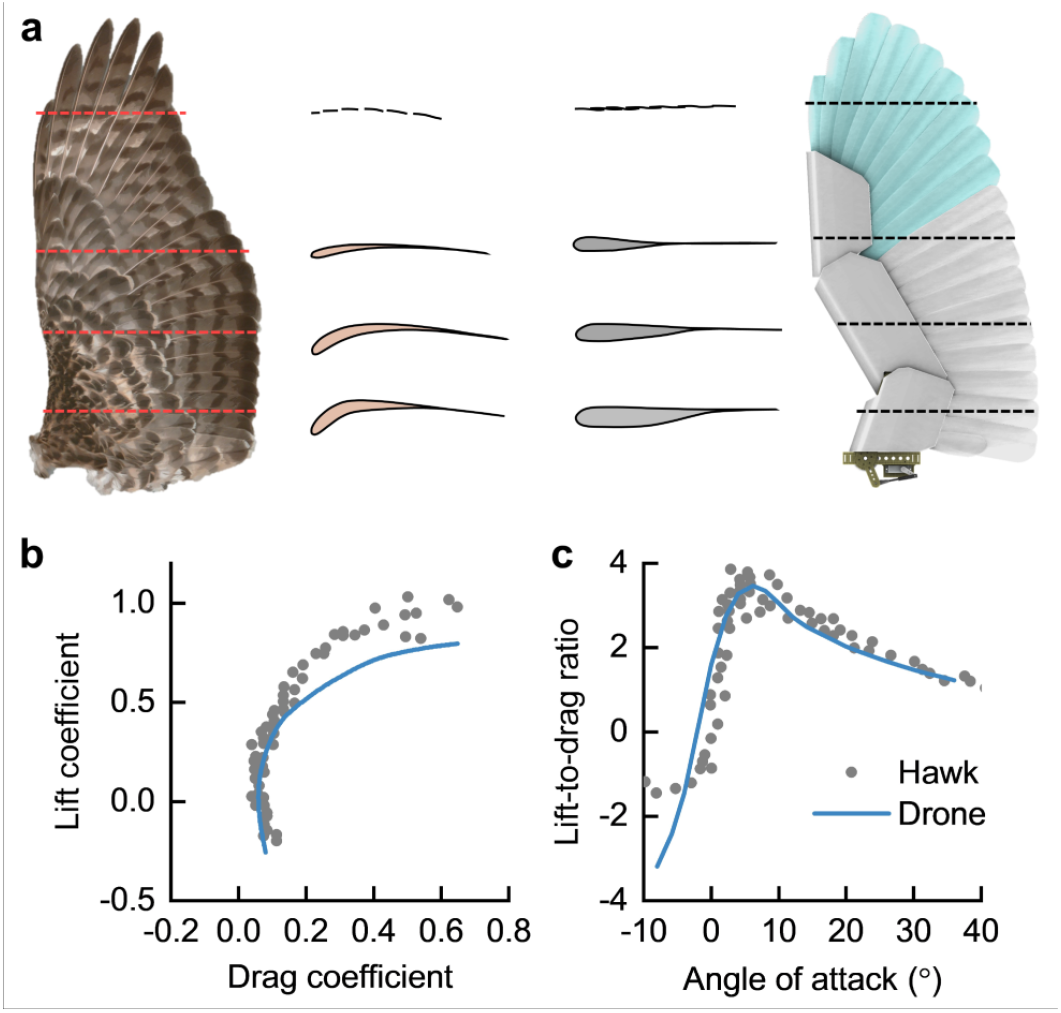
Wing airfoils, and lift and drag generation of the mechanical feathered wing and a hawk wing. **a**, Wing airfoils in a hawk wing (adapted from^50^) and the mechanical wing. **b**, Lift coefficient as a function of drag coefficient. **c**, Lift-to-drag ratio as a function of the angle of attack. Data of bird wing were reproduced from^45^ with permission.

**Extended Data Fig. 5.**
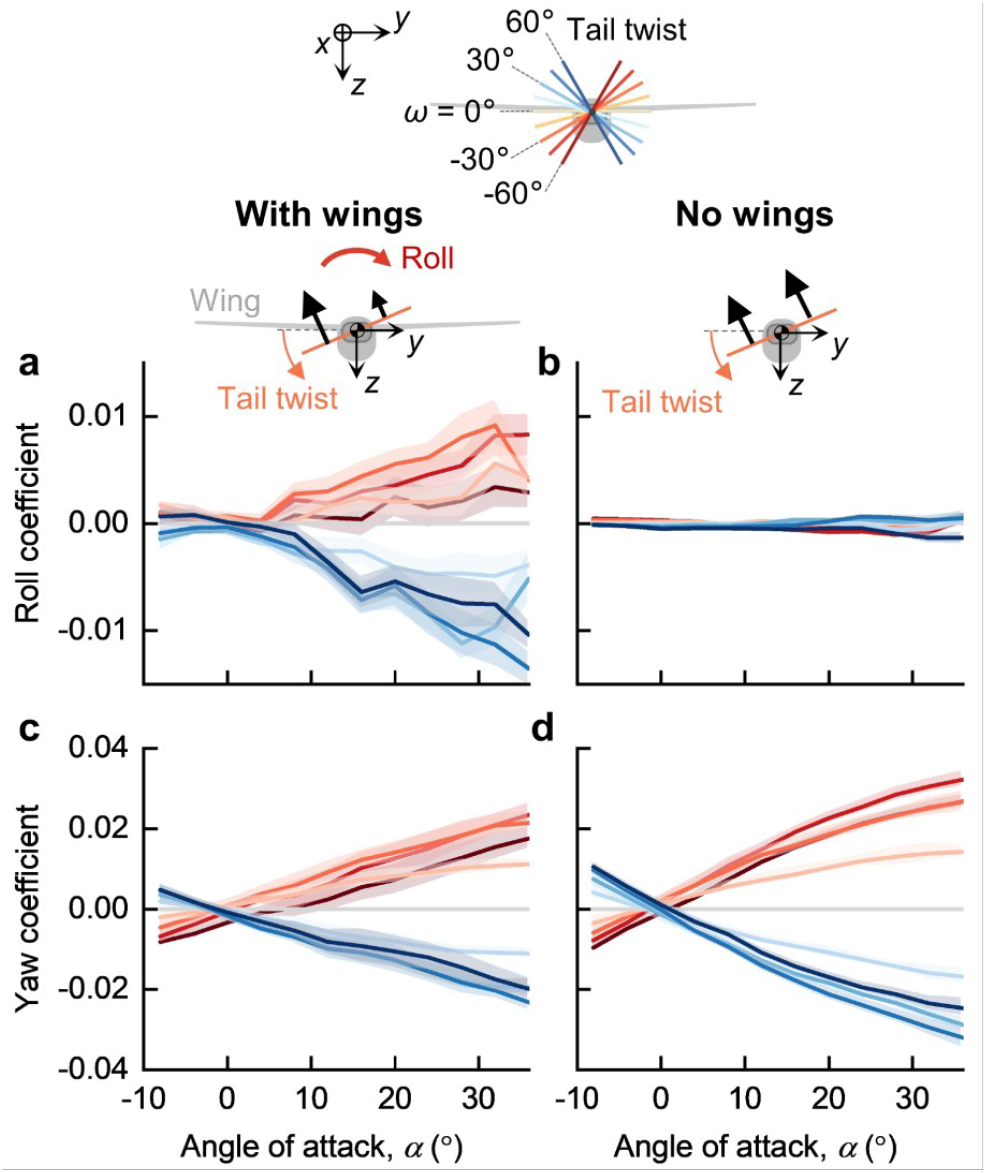
Generation of roll and yaw moments by tail twist with the CG located at the tail rotational axis. **a**,**b**, Roll coefficients with and without presence of the wings. **c**,**d**, Yaw coefficients with and without presence of the wings.

**Extended Data Fig. 6.**
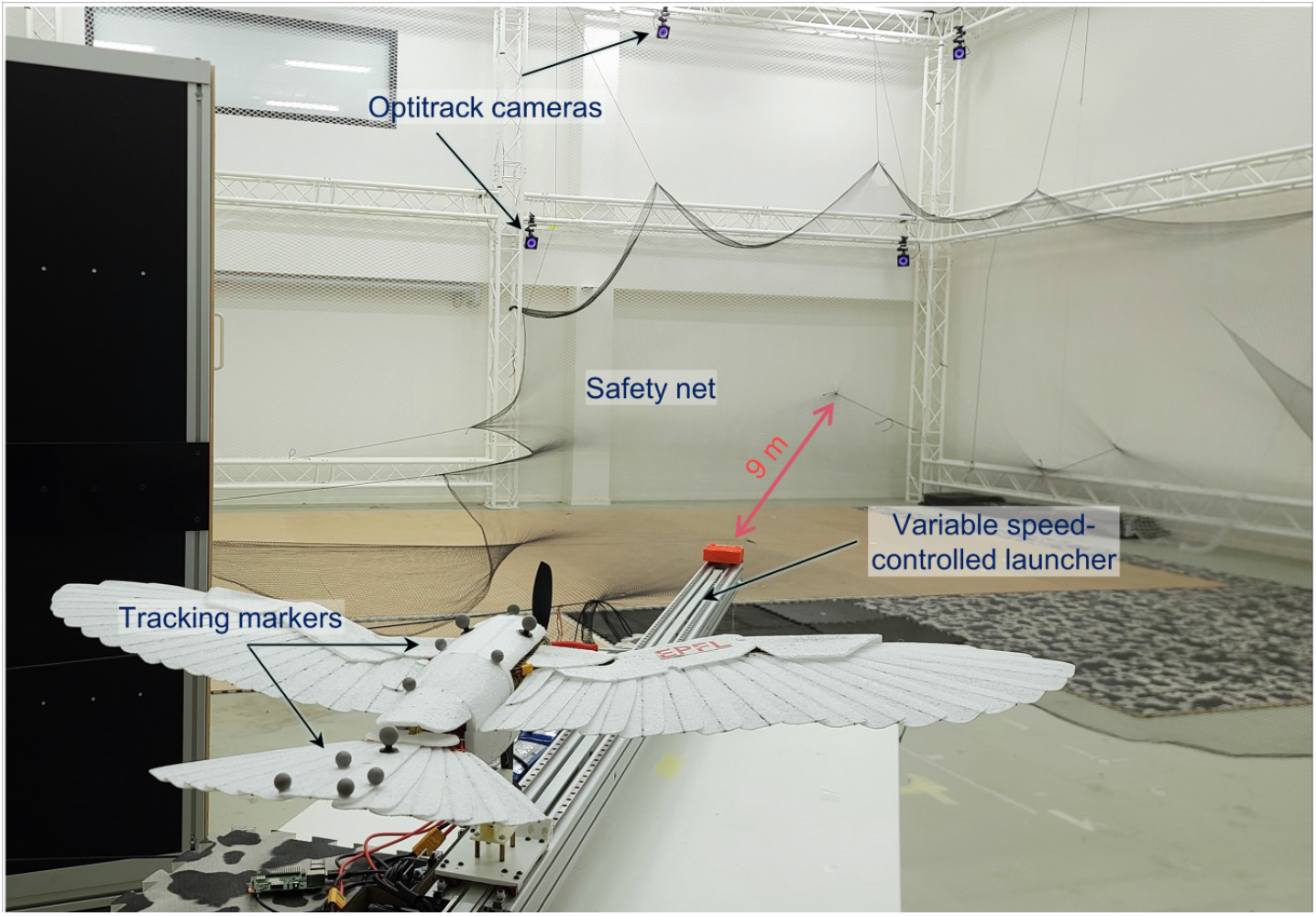
Indoor flight setup. We conducted flight tests in a 10 m × 10 m × 8 m indoor flight arena equipped with a safety net and a motion tracking system (26 Optitrack cameras, 240 Hz). The drone was launched by a custom-built, programmable linear launcher.

**Extended Data Fig. 7.**
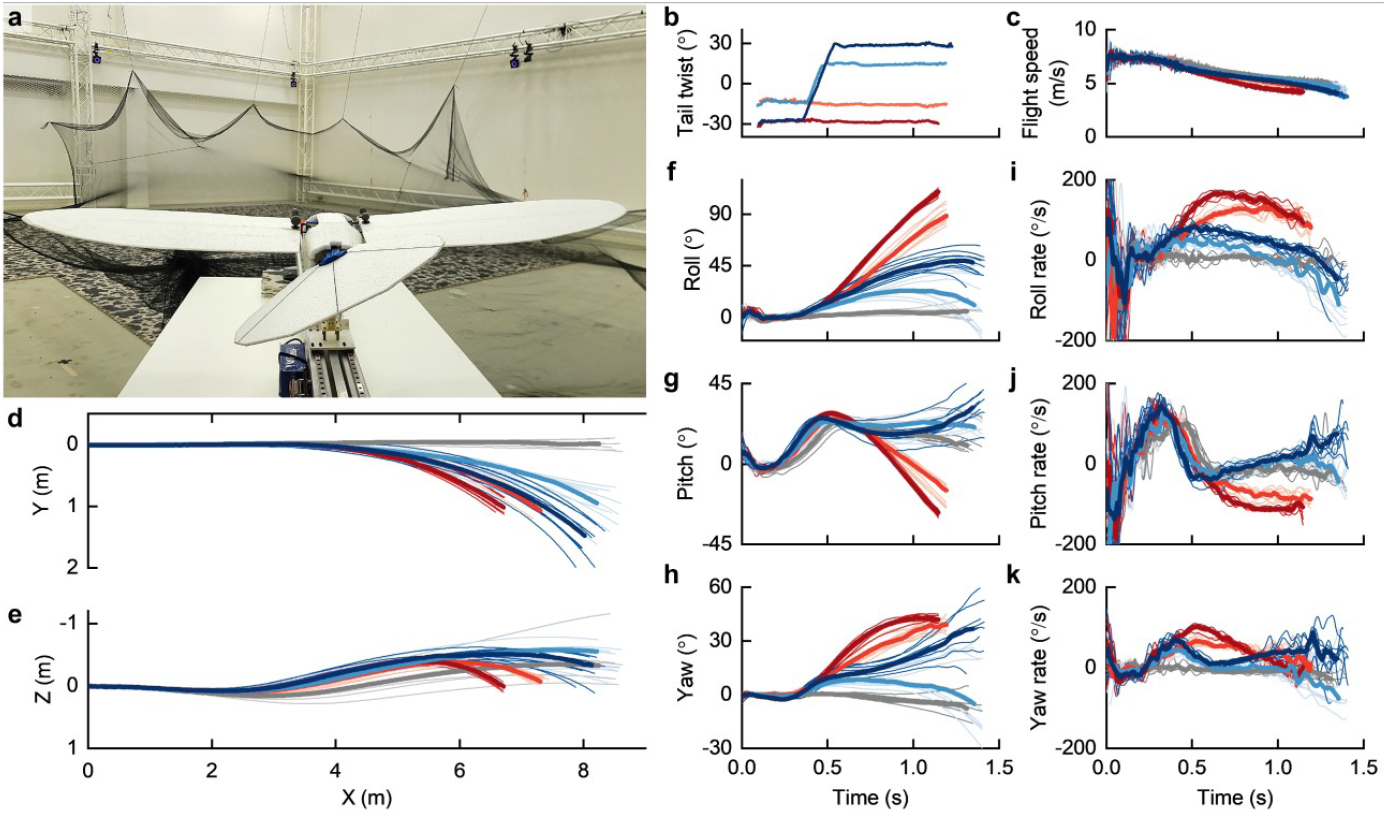
Steady banked turns with a rudderless fixed-wing drone. **a**, The rudderless fixed-wing drone can twist its tail around the body axis. **b**, Tail-twist angles in four flight patterns: banked turns with the tail twists of 15° and 30° without (red) and with (blue) reversal at the end of the turn. **c**, Flight speeds during turning. **d**,**e**, Flight trajectory in the *XY*-plane (**d**) and the *XZ*-plane (**e**). **f**–**h**, Body attitude angles: roll (**f**), pitch (**g**), and yaw (**h**). **i**–**k**, Body rates: roll rate (**i**), pitch rate (**j**), and yaw rate (**k**). In **d** to **k**, thin lines represent individual flight trials; thick lines are the mean, respectively. Data in grey color represent trimmed flights.

**Extended Data Fig. 8.**
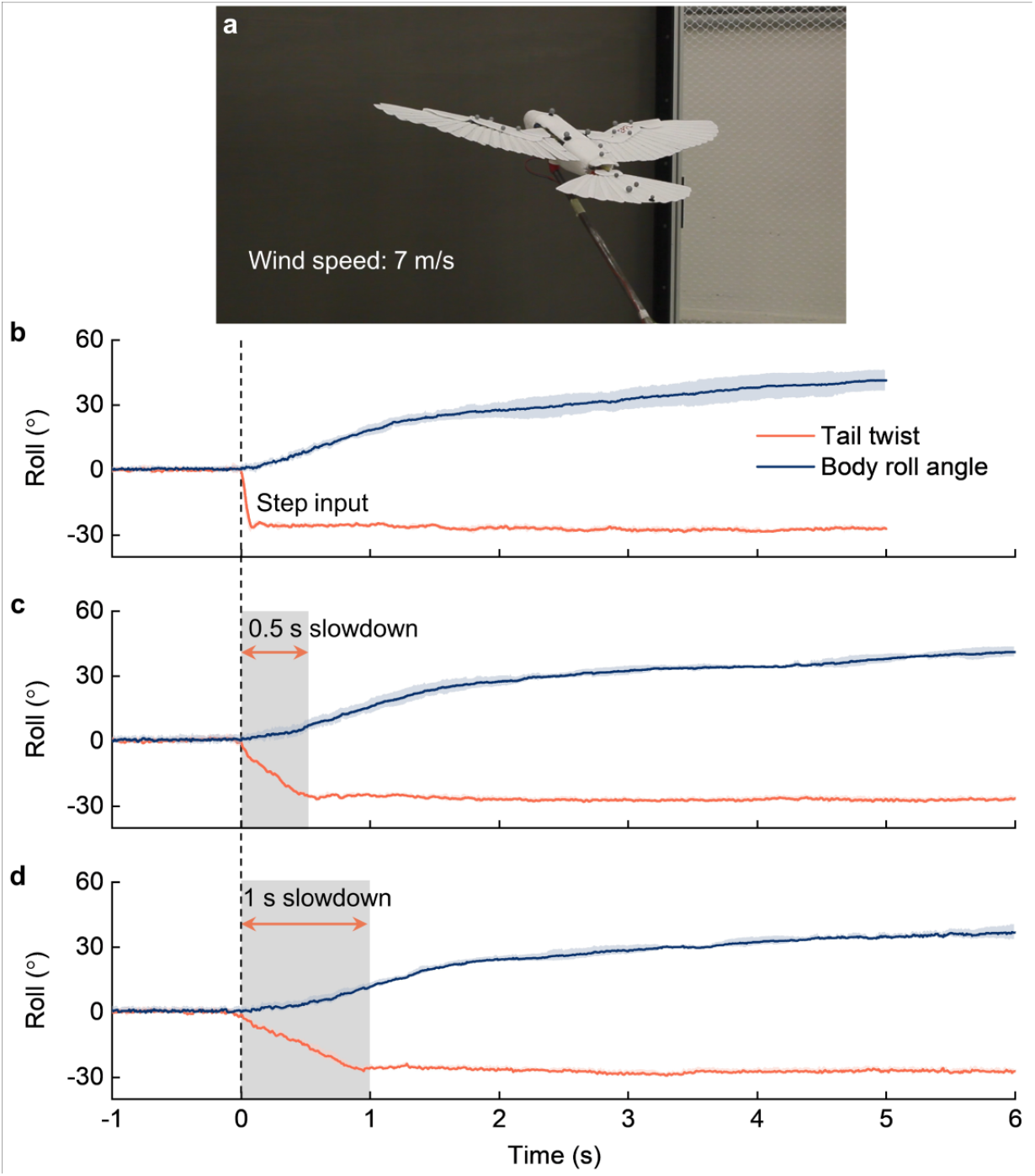
Roll motion is minimally affected by the twisting rate of the tail. **a**, Experimental setup. The drone, capable of 1 degree-of-freedom rolling around its body axis, was commanded to twist its tail from 0 to - 30° with step input, and 0.5 s and 1.0 s slowdown inputs. The drone’s angle of attack was set at 20°, where generated lift is sufficient to compensate for the weight of the drone. **b-d**, Roll responses with various twisting rates: step input (**b**), 0.5 s slowdown (**c**), and 1.0 s slowdown (**d**).

**Extended Data Fig. 9.**
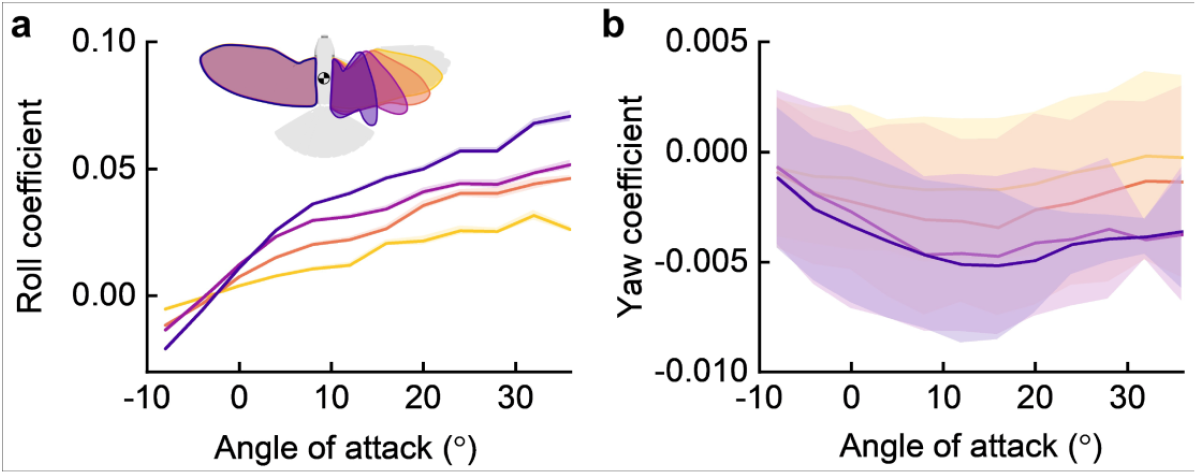
Roll and yaw moments generated by wing shape morphing. **a**,**b**, Wing shape morphing effectively generates roll moment (**a**) but also causes adverse yaw (**b**).

**Extended Data Fig. 10.**
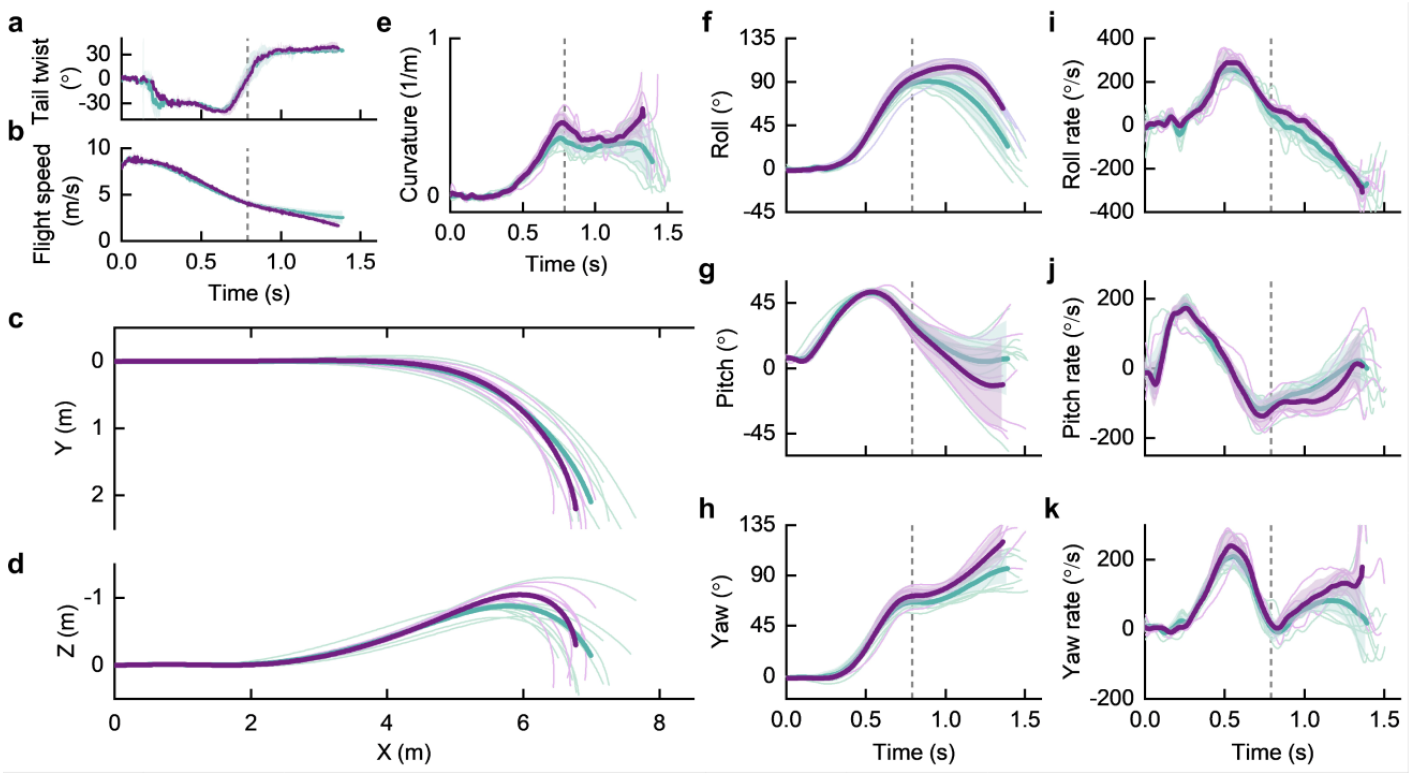
Sharp right turns by tail twist with the coordination of wing morphing in the exit phase to return the drone to the level flight. Green and violet denote the turns with the left wing folding triggered at the peak of the altitude (n =10) and after 0.3 second delay (n =7), respectively. Thin lines represent individual flight trials; thick lines and shaded areas are the mean and standard deviation from the mean, respectively. **a**,**b**, Tail twist angles (**a**) and flight speeds (**b**). **c**,**d**, Flight trajectories in the *XY*-plane (**c**) and the *XZ*-plane (**d**). **e**, Curvature of the turns. **f**–**h**, Body attitude angles: roll (**f**), pitch (**g**), and yaw (**h**). **i**–**k**, Body rates: roll rate (**i**), pitch rate (**j**), and yaw rate (**k**).

**Extended Data Fig. 11.**
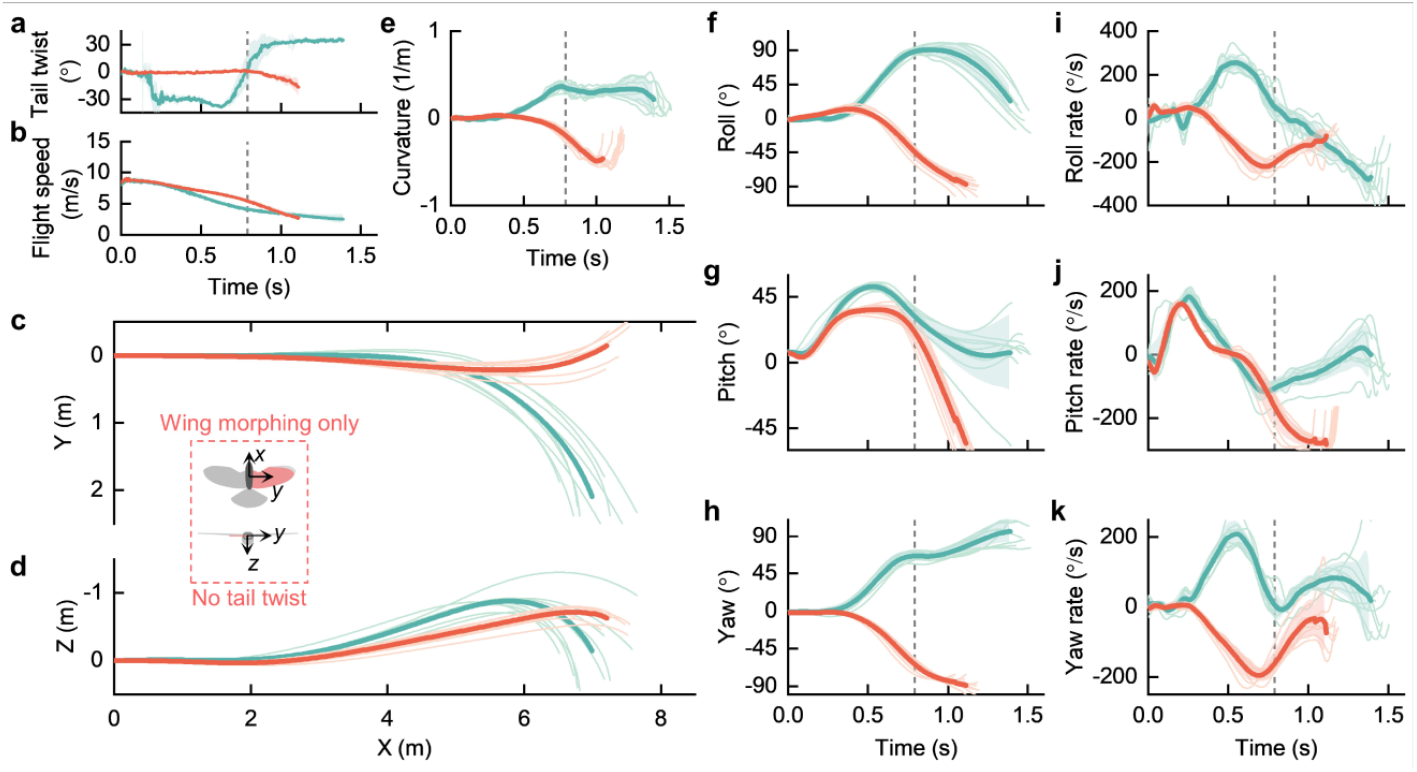
Banked turns by tail twist (green) and asymmetric wing morphing only (red). **a**,**b**, Tail twist angles (**a**) and flight speeds (**b**). **c**,**d**, Flight trajectories in the *XY*-plane (**c**) and the *XZ*-plane (**d**). **e**, Curvature of the turns. **f**–**h**, Body attitude angles: roll (**f**), pitch (**g**), and yaw (**h**). **i**–**k**, Body rates: roll rate (**i**), pitch rate (**j**), and yaw rate (**k**). Asymmetric wing morphing induces adverse yaw that causes the drone to turn in the opposite direction.

**Extended Data Table 1.** Design parameters of the drone.

| **Parameter** | **Value** | **Mass contribution (%)** |
| --- | --- | --- |
| Total mass | 460 g | 100 |
| Wingspan | 1.1 m |  |
| Body length | 0.54 m |  |
| One wing mass | 59.4 g | 12.9 |
| Wing length | 0.5 m |  |
| Wing mean chord | 0.191 m |  |
| Humerus length | 0.09 m |  |
| Humerus angle range, $\theta_1$ (min - max) | 28.5° – 73.5° | |
| Ulna length | 0.13 m |  |
| Ulna angle range, $\theta_2$ (min - max) | 46.7° – 115.4° | |
| Manus length | 0.1 m |  |
| Manus angle range, $\theta_3$ (min - max) | 25.2° – 128.1° | |
| Tail mass | 14 g | 3 |
| Tail length | 0.25 m |  |
| Maximum tail spread angle | 120° |  |

## Notes

### Competing Interest Statement

The authors have declared no competing interest.

